# Layer V Neocortical Neurons From Individuals With Drug-Resistant Epilepsy Show Multiple Synaptic Alterations but Lack Somatic Hyperexcitability

**DOI:** 10.1101/2025.02.26.640307

**Authors:** Luis A. Márquez, Christopher Martínez-Aguirre, Estefanía Gutierrez-Castañeda, Ernesto Griego, Isabel Sollozo-Dupont, Félix López-Preza, Mario Alonso-Vanegas, Luisa Rocha, Emilio J. Galván

## Abstract

Although neuronal hyperexcitability is the primary mechanism underlying seizure activity in epilepsy, little is known about how different neuronal mechanisms at different organizational levels contribute to network hyperexcitability in the human epileptic brain. In this study, we determined a series of cellular and synaptic properties of layer V pyramidal neurons from neocortical tissue of patients with drug-resistant epilepsy that may contribute the hyperexcitable state associated with epilepsy. Using the whole cell, patch-clamp technique, and extracellular recordings, we determined the passive and active electrophysiological properties of layer V pyramidal neurons with regular spiking phenotypes from temporal, parietal, and frontal neocortices surgically resected from individuals with drug-resistant epilepsy. Also, the glutamatergic strength, the synchronicity between presynaptic volleys and field excitatory postsynaptic potentials, and short-term, frequency-dependent plasticity were determined at the synaptic level. Lastly, biocytin-filled pyramidal neurons were used to perform post hoc digital reconstructions and morphometric analyses. The collected data revealed that pyramidal neurons exhibit minimal spontaneous activity, similar resting membrane potentials, and input resistance values among the temporal, parietal, and frontal neocortices. Although frontal neurons were more hyperexcitable than temporal and parietal neurons, the firing output was comparable to that previously observed in non-pathological human tissue. The digital reconstructions confirmed the identity of pyramidal neurons and revealed alterations in dendritic complexity. In contrast, the analyses of the extracellular recordings uncovered significant desynchronization between presynaptic excitability and postsynaptic activity and loss of short-term depression in response to repetitive stimulation within the gamma range (30 Hz). Our data suggest that neocortical layer V pyramidal neurons from individuals with drug-resistant epilepsy are not necessarily hyperexcitable at the somatic level. Instead, synaptic alterations, such as synaptic desynchronization and a loss of frequency-dependent short-term depression may significantly contribute to the hyperexcitable state observed during seizure activity.

## Introduction

Although significant progress has been made in animal models to understand the intrinsic and synaptic alterations that can trigger neuronal hyperexcitability —a hallmark of epilepsy alongside recurrent seizure activity [1–6]— the translation of these neurophysiological findings to the brain of patients with epilepsy remains largely unverified, with few exceptions [7,8]. In this line of work, neocortical hyperexcitability at network level is recognized as a central mechanism underlying seizure activity [7,9]. However, the potential contribution of cellular mechanisms at lower organizational levels, ranging from cellular to synaptic, is less understood [3,10,11].

A more realistic approach to studying the cellular and synaptic mechanisms underlying the hyperexcitable state associated with epilepsy is to directly examine pathological brain tissue from surgical procedures in epileptic patients. Nevertheless, given the limited access and the bioethical concerns in obtaining human tissue, these findings are regularly compared to results from analogous structures in the rodent brain. Furthermore, a growing number of studies have documented numerous differences between rodent and human brain structures. For example, the process of synaptic integration in pyramidal cells (PCs) of the neocortex differs between rodents and humans [12–14]. These subtle differences extend to the functional expression of ion channels, their biophysical properties, neuronal firing, short-term plasticity, and efficiency in information processing [15,16]. Likewise, morphological differences exist between the human and rodent neocortex, including dendritic complexity of pyramidal neurons [17,18]. These differences, along with the limited understanding of the intrinsic properties and synaptic physiology of human neurons resembling their physiological state [10,11], may potentially lead to misinterpretations of the cellular mechanisms underlying the hyperexcitable state of epilepsy.

Given that layer V PCs have been identified as a primary source of interictal discharges in epilepsy [19–21] and exhibit unique morphological and biophysical characteristics within the laminar structure of the human neocortex [22,23], this study examined the biophysical, synaptic, and morphological differences of layer V PCs in the temporal, parietal, and frontal cortices surgically resected from patients with drug-resistant epilepsy (DRE). Patch-clamp recordings in acute human neocortical slices were used to classify PCs based on their membrane properties and firing patterns, while post hoc reconstructions revealed anatomic details of layer V PCs. Extracellular recordings assessed synaptic strength, synchronization between presynaptic volleys and excitatory postsynaptic potentials, and short-term plasticity in response to gamma-range stimulation. Due to the inherent challenge of obtaining ‘healthy’ tissue for comparative purposes, see [11], the findings obtained in this study were interpreted alongside studies using non-pathological tissue. Our results suggest that intrinsic cellular and synaptic mechanisms in layer V PCs may differentially contribute to the dominant hyperexcitable state during seizures in patients with DRE.

## Material and methods

### Preparation of Human Cortical Slices

Neocortical brain tissue of the temporal, parietal, or frontal neocortex was obtained during surgical procedures of 10 patients with DRE (Table 1). Tissue samples were collected between June 21, 2022, and July 23, 2023. Immediately after resection, the epileptic neocortex was placed in ice-cold isotonic buffer solution (in mM: 320 sucrose, 1 EDTA, 5 Tris-HCl; pH = 7.4) with continuous bubbling of carbogen (95% O_2_/5% CO_2_ at 0.5 L/min). The transportation time between the resection and the beginning of slice preparation in the neurophysiology laboratory was less than 45 minutes. The experimental procedures (protocols: 055/2018 and 055/2019) were approved by our institution’s internal ethics committee. The procedures for obtaining brain tissue from epileptic patients (protocol FR-2019-785-008) were approved by the internal ethics committees of the involved medical institutions, in accordance with the Declaration of Helsinki. Written informed consent was obtained from all patients. Patients’ clinical variables, such as sex, age, epileptic focus location, and drug treatment, are summarized in Table 1.

**Table 1.** Clinical variables of patients with drug-resistant epilepsy.

| ID | Sex | Age (years) | Age of seizure onset (years) | Duration of epilepsy (years) | Frequency of the seizures (per month) | Location of the focus | ASMs administrated before surgery |
| --- | --- | --- | --- | --- | --- | --- | --- |
| HUM-209 | F | 21 | 10 | 11 | 7 | Left temporal lobe | LEV, OXC, NMP |
| HUM-210 | F | 30 | 13 | 17 | 20 | Right frontal lobe | OXC, LEV, TOP |
| HUM-211 | M | 21 | 13 | 8 | 12 | Right temporal lobe | LEV, VPA, PAL, OXC, ZSM |
| HUM-214 | M | 2.6 | 1.2 | 1.4 | 10 | Right frontal lobe | LEV, LAC |
| HUM-221 | F | 27 | NA | NA | NA | Right temporal lobe | LEV, LAC, PAL, OXC |
| HUM-215 | M | 16 | 10 | 6 | 150 | Left frontal lobe | LAC, LEV, CLB, CZM, VAP, LAM |
| HUM-216 | F | 2 | 0.9 | 1.1 | 3 | Left parietal lobe | LEV, CLB, VAP, LAM, OXC |
| HUM-217 | F | 7 | 1.25 | 5.75 | NA | Left parietal lobe | LEV, OXC, BRI, PAL, LAC, VPA, CLB |
| HUM-218 | F | 24 | 16 | 8 | NA | Right frontal lobe | LAM, VAP, LEV |
| HUM-219 | M | 4 | 2.75 | 1.25 | 1 | Right parietal lobe | VPA |

ASMs, Antiseizure medications; BRI, brivaracetam; CBZ, carbamazepine; CLB, clobazam; CZM, clonazepam; F, female; LAC, lacosamide; LAM, lamotrigine; LEV, levetiracetam; M, male; NA, not available; NMP, nimodipine; OXC, oxcarbazepine; PAL, perampanel; TOP, topiramate; VPA, valproic acid; ZSM, zonisamide

In the laboratory, brain tissue samples were transferred to a frosty sucrose solution, maintaining the carbogen bubbling. The sucrose solution composition was (in mM): 210 sucrose, 2.8 KCl, 2 MgSO_4_, 1.25 Na_2_HPO_4_, 25 NaHCO_3_, 1 MgCl_2_, 1 CaCl_2_, and 10 D-glucose. Then, coronal slices of the human neocortex (350 µM thickness) were obtained from neocortical tissue blocks dissected from the tissue samples. The neocortical tissue blocks were sliced perpendicular to the pial surface using a vibratome (Leica VT1000S; Nussloch, Germany). Next, the acute slices were maintained at 34°C for 25–30 minutes in artificial cerebrospinal fluid solution (ACSF; pH ≈ 7.30–7.35) with the following composition (in mM): 125 NaCl, 2.5 KCl, 1.25 Na_2_HPO_4_, 25 NaHCO_3_, 4 MgCl_2_, 1 CaCl_2_, and 10 D-glucose. Then, the slices were maintained at room temperature for at least 1 hour before the whole-cell patch clamp and extracellular recordings were performed. For the recordings, an extracellular solution (modified ACSF) was continuously perfused to slices containing (in mM): 125 NaCl, 2.5 KCl, 1.25 Na_2_HPO_4_, 25 NaHCO_3_, 1.5 MgCl_2_, 2.5 CaCl_2_, and 10 D-glucose. All recordings were performed at 32.5 ± 1°C.

### Whole-Cell Recordings

The PCs were visualized with infrared differential interference contrast optics coupled to an FN1 Eclipse microscope (Nikon Corporation, Minato, Tokyo, Japan). PCs located in layer V were identified based on their characteristic shape (broad base, pointed apex, and a single dendritic elongation pointing towards the pia-cortex) and position. The patch pipettes were pulled from borosilicate glass using a micropipette puller (P97, Sutter Instruments, Novato, CA, USA). The pipette tips had a resistance of 4–6 MΩ when filled with an intracellular solution (pH ≈ 7.20–7.28) with the following composition (in mM): 135 K^+^-gluconate, 10 KCl, 5 NaCl, 1 ethylene glycol-bis(β-aminoethyl ether)-N,N,N′,N′-tetraacetic acid (EGTA), 10 N-(2-hydroxyethyl)piperazine-N′-(2-ethane sulfonic acid) (HEPES), 2 Mg^2+^-ATP, 0.4 Na^+^-GTP, and 10 phosphocreatine. Biocytin (0.04%) was routinely added to the intracellular solution for post hoc digital reconstructions and morphological analysis of the recorded neurons. Whole-cell patch-clamp recordings were performed using an Axopatch 200B amplifier (Molecular Devices, San José, CA, USA), digitized at 40 kHz, and filtered at 1 kHz with a Digidata 1550B (Axon Instruments, Palo Alto, CA, USA). Digital signals were acquired and analyzed offline with the pCLAMP 11.2 software (Molecular Devices).

### Determination of Passive and Active Electrophysiological Properties

The resting membrane potential was measured after the initial break-in from giga-seal to whole-cell configuration in the current-clamp configuration mode, followed by determining the synaptic spontaneous activity, monitored for 3 minutes in the gap-free mode. Next, a series of hyperpolarizing and depolarizing current pulses (from -300 pA to 60 pA; 30 pA steps, 1 s of duration) were somatically injected to determine the current-voltage (I–V) relationship, somatic input resistance, membrane time constant, and membrane capacitance. Input resistance was determined as the slope of a linear fit (*f*(*x*) = *mx* + *b*) to the steady-stage I–V plot elicited by subthreshold current injection (-60 to 0 pA). The time constant was calculated by fitting a single exponential function 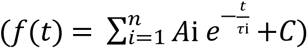 to a voltage response elicited by a current pulse of -30 pA. Membrane capacitance (pF) was calculated as time constant to input resistance ratio. Rheobase and latency to first action potential (AP) were calculated using depolarizing current ramps (30 pA steps, 500 ms duration). The AP kinetic measurements included half-width, maximal depolarization slope, and maximal repolarization slope. These kinetic parameters were depicted with phase plots, constructed by plotting the first derivative of membrane potential *dV/dt* (mV/ms) with the membrane potential. To determine the recorded cells’ excitability level, a series of firing rate–current curves were constructed by computing the number of elicited APs in response to increasing current injection from 0 to 390 pA (30 pA steps; 1s duration). From these curves, the neuronal gain was calculated as the slope of a linear fit to the steady-stage rate firing–current plot obtained by current injection from 90 to 210 pA.

### Classification of Neocortical Cells According to Firing Pattern

PCs’ electrophysiological identity was determined by visualizing neuronal firing patterns and analyzing spike frequency adaptation [24]. The spike frequency adaptation was calculated as the slope of the linear fit of the relationship between the interspike interval (ms) and each interval’s latency (ms), elicited with a current pulse of 210 pA (1 s). The slope values revealed that PCs had three different firing patterns: regular spiking, intrinsic burst, and adaptative firing. In addition, the instantaneous firing frequency was computed as the reciprocal value of the interspike interval during consecutive APs. These data were plotted as the instantaneous firing frequency of each AP elicited by increasing current pulses (120 to 360 pA; 1 s) for each neuronal firing type.

### Extracellular Recordings

Extracellular recordings were used to analyze the synaptic strength. A bipolar stimulation electrode was placed in layers I and II (LI/II), and the resulting field excitatory postsynaptic potential (fEPSP) was recorded in the dendritic region of layer V with a borosilicate pipette (resistance: 1–2 MΩ when filled with 3M NaCl solution). The current pulses were delivered via a high-voltage isolation unit (A365D; World Precision Instruments, Sarasota, FL, USA) under the command of a Master-8 pulse generator (AMPI, Jerusalem, Israel). Electrical responses were amplified using a Dagan BVC-700A amplifier (Minneapolis, MN, USA) connected with a 100x gain headstage (Dagan, model 8024) and a high-pass filter set at 0.3 Hz. Additional electrical noise suppression was achieved using a Humbug device (Quest Scientific Instruments; North Vancouver, BC, Canada). The evoked fEPSPs were displayed on a computer-based oscilloscope and digitalized with an A/D converter (BNC-2110) for storage and offline analysis with LabVIEW 7.1 software (National Instruments, Austin, TX, USA).

### Characterization of fEPSP Waveform, Paired-Pulse Ratio, Frequency-Dependent Short-Term Plasticity and neuronal synchronization

The fEPSPs were acquired at 0.067 Hz with current pulses (200 to 250 µA) of 100 µs duration. Ten consecutive sweeps were averaged and then analyzed for measurements of the kinetic parameters of the fEPSP waveform. For each fEPSP, the amplitude (measured in mV), half-width (ms), peak time (ms), fiber volley (FV) amplitude (mV), and slope (mV/ms from 10% to 80% of response) were measured with custom made software written in LabVIEW 7.1 or pCLAMP 11.2 software.

For paired-pulse ratio (PPR) analysis, 12 consecutive synaptic responses with paired stimulation (inter-stimulus interval of 60 ms; 15%–20% of maximal response) were acquired at 0.067 Hz. The PPR was calculated as the arithmetic division between the amplitude of the second response and the amplitude of the first response. To determine short-term plasticity in response to brief high-frequency stimulation, a baseline response of fEPSP was configured at 50% of its maximal amplitude and then acquired for 10 min, as previously reported [25]. Then, a train of 10 current pulses (100 µs of duration) at 30 Hz was delivered in LI/II. The train’s evoked synaptic responses were normalized to the slope of its first synaptic response and plotted as the number of stimuli vs. normalized fEPSP slope (%). Additionally, spontaneous synaptic events were detected and quantified during train stimulation. These events occurred more than 10 ms after electrical stimuli and were therefore considered spontaneous, likely generated by short-term synaptic properties rather than evoked activity. The spontaneous events were detected using pCLAMP 11.2 software based on the following criteria: an inward-directed waveform, amplitude > 0.15 mV and duration > 1 ms.

To establish the relationship between presynaptic action potentials (fiber volleys, FV) and fEPSPs by neocortical region, a Pearson correlation analysis (*r*^2^) was used, and the least squares method was applied to find the equation of best-fitting curve or line to the dataset considering the next formula: *y* = *mx* + *b*, where *y* is the dependent variable (fEPSP), *x* is the independent variable (FV), *m* is the slope of the line, and *b* is the *y*-intercept. As a reference group for normal synaptic synchronization, we performed a similar analysis using previously published data from layer V of the rat prefrontal cortex [26]. The formulas to calculate the slope (*m*) and intercept (*b*) of the line are derived from the following equations:

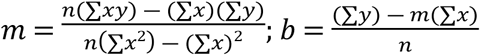, where n is the number of data points, ∑*xy* is the sum of the product of each pair of *x* and *y* values, ∑*x* is the sum of all *x* values, ∑*y* is the sum of all y values, and ∑*x*^2^ is the sum of the squares of *x* values.

The observed values of *x* were used in conjunction with the formula *y* = *mx* + *b* to calculate the predicted values of *y*. The residual values were estimated as the difference between the observed values of *y* and the predicted *y* values. Lastly, an exploratory analysis (EA) of the residuals was conducted to assess the feasibility of using the least squares method. The EA involved the Kolmogorov–Smirnov test, which assesses whether the residuals follow a specific distribution, helping us evaluate the normality assumption required for the least squares method. Additionally, the biases of the residuals were determined using a one-sample Student’s t-test. This test allowed us to evaluate whether the mean of the residuals significantly deviated from zero, helping to identify any systematic bias in the model’s predictions. A non-significant result would suggest that the model’s predictions were unbiased.

### Immunostaining of Biocytin-Filled Neurons and Confocal Image Acquisition

Slices containing the biocytin-filled neurons obtained from the patch-clamp recordings were fixed in 4% PFA solution in 0.1 M PB (pH = 7.4) and kept at 4°C for 24–48 hours for fixation. Then, the slices were washed with PBS and treated with a blocking/permeation buffer (PBS containing 3% BSA and 0.3% Triton X-100) at room temperature for 2 hours. Subsequently, the slices were exposed to Streptavidin Texas Red (diluted at 1:500, Vector Laboratories; SA-5006) overnight at room temperature. Finally, the sections were washed again and mounted using a Vectashield Vibrance curing antifade mounting medium (Vector Laboratories; H-1800). Photomicrographs were acquired using confocal microscopy (LSM 800 with Airyscan; Carl Zeiss) [27]. The excitation wavelength for Texas Red was 561 nm. Multiple tile z-stack series were captured from each cell and reconstructed using Zen 2.3 (Carl Zeiss) and ImageJ (NIH Image) software.

### Analysis of Dendritic Complexity

Dendritic arborization was quantified by counting the numbers of total and primary dendrites, the total dendritic length, the axon extension, and the total projection area, considered as the area occupied by all dendrite segments for each neuron, using ImageJ software coupled with the Sholl analysis plug-in v3.4.2 for morphometric analysis [28,29]. Primary neurites were defined as those that originated directly from the soma. Sholl analysis was used to determine the complexity of dendritic branching. Concentric circles were drawn every 10 μm from the soma, and the number of intersections between the increasing circles and the dendritic arbor was quantified, as well as the number of branch points and end tips.

### Data Analysis

During experimental procedures, the experimenters were blinded to both the tissue brain region origin and the clinical variables of the patients. All data are numerically expressed as mean ± SEM unless otherwise stated. The violin plots show the median and the first and third quartiles. The normal distribution of data was validated with the Kolmogorov– Smirnov test (*p* = 0.05). No outliers were removed from the data. The group size per cortex type is expressed as the number of cells or slices obtained from *n* patients. For the morphometric analyses, descriptive statistics were performed for each parameter, including the numbers of total and primary dendrites, total dendritic length, axon extension, and total projection area. The comparability among experimental conditions was assessed by a ratio-paired Student’s t-test, one-way analysis of variance (ANOVA), or mixed-effects ANOVA, as appropriate. Tukey’s post hoc test was computed for multiple comparisons among experimental groups only when F achieved minimal statistical significance. For all the experiments, data were considered significant if *p* < 0.05.

## Results

### Identification of Neocortical Pyramidal Cells by Firing Pattern From Patients With Drug-Resistant Epilepsy

We recorded 35 PCs in acute slices of the human neocortex (Fig 1A) using the whole-cell configuration of the patch-clamp technique; the intracellular solution routinely included biocytin to perform intracellular labeling. The tissue was surgically resected from 10 patients with DRE ranging from 2 to 30 years old of both sexes (Fig 1B, upper panel and Table 1). The average time between the surgical resection, brain slice preparation, and slice placement in the recording chambers was 4 ± 0.5 h. According to its origin, the tissue was obtained from the temporal, parietal, and frontal neocortices (Figs 1A and 1B, middle panel).

**Fig 1.**
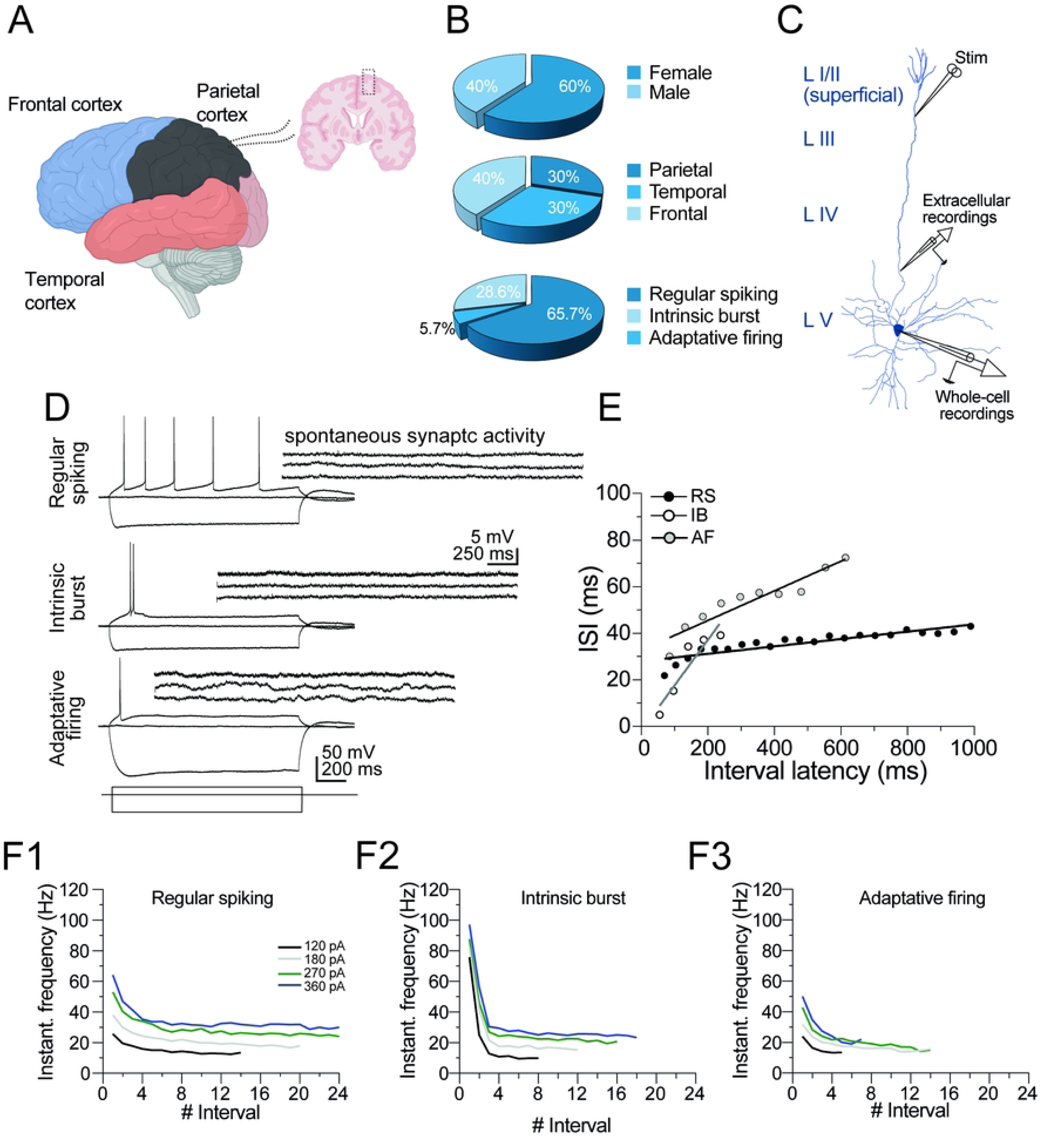
Layer V pyramidal cell types from the human neocortex with drug-resistant epilepsy. **(A)** Schematic representation showing the cortical sections used for this study. Acute cortical slices were prepared from temporal, parietal, and frontal cortices resected from drug-resistant epileptic brains. **(B)** Pie charts with the percental distribution of the examined brain tissue (n = 10 patients) by sex (top panel), cortex type (middle panel), and neuronal firing type (lower panel, n = 35 cells). **(C)** Schematic representation for the whole-cell patch clamp and extracellular recordings in PCs of human cortical slices. The patch pipette was placed in layer V PCs for whole-cell recordings. For extracellular recordings, the stimulation electrode was placed in the superficial layer I/II, and the synaptic responses were obtained in the deep layer V. **(D)** Representative examples of whole-cell recordings showing the firing patterns recorded from PCs. The upper left panel shows a regular spiking (RS) neuron; the middle panel shows an intrinsic bursting (IB) neuron, and the lower panel shows the adaptative firing (AF) neuron. A common characteristic of the neurons in this study was their low spontaneous synaptic activity obtained at each neuron’s resting membrane potential (notice the calibration bar). Neurons with depolarized resting membrane potentials usually exhibited robust spontaneous synaptic activity. Those cells were ruled out from this study. **(E)** Analysis of neuronal firing adaptation. A representative graph of the interval latency of each AP vs. inter-spike interval (ISI, ms). This analysis corroborated the differential nature between the RS, IB, and AF neurons. Notice that the decreased ISI in AF neurons is higher than in IB and RS neurons. **(F)** The IFF trajectories of RS, IB, and AF neurons (F1–F3, respectively) in response to different current magnitudes.

The somata of PCs were positioned in the middle region of layer V of the neocortex (Fig 1C), 50–150 µm below the slice surface. Under DIC microscopy, the PCs somata size was 22.2 ± 4.3 µm, and visibly different from local interneurons that exhibited smaller rounded or bifurcated somata. Strictly pyramidal cell-like morphology, including a broad base, a pointed apex, and the protrusion of at least one major dendritic branch pointing towards the cortex-pia, was an inclusion criterion for patch-clamp recordings. Following the initial membrane break-in from giga-seal to whole-cell configuration, we switched to the current-clamp mode, and the spontaneous synaptic activity (SSA) was monitored at the resting membrane potential (RMP) of the recorded cells. Since the tissue was obtained from epileptic individuals, the SSA was a parameter routinely used to identify possible hyperexcitable cells. However, PCs included in this study barely exhibited SSA at their RMP (right insets Fig 1D).

Next, we classified the PCs according to their firing pattern (Fig 1D). Independently of the origin of the epileptic tissue, we found that 65.7% of PCs exhibited a regular spiking firing pattern (RS) phenotype characterized by stable AP firing, weak AP adaptation, and stable AP amplitude and afterhyperpolarization; 28.6% of PCs exhibited an intrinsic bursting (IB) firing pattern characterized by clustered spikes (2–3) observed during depolarizing plateau potentials and followed by quiescent periods; the remaining 5.7% of PCs exhibited adaptative firing (AF) in which one AP with a defined afterhyperpolarization and strong adaptation was recorded (Fig 1D, lower panel). Since the cells in this study did not exhibit SSA, we assumed that the firing patterns were intrinsically regulated and not derived from synaptic activity impinging on the recorded PCs, as summarized in the voltage traces and the accompanying SSA in Fig 1D.

Next, to corroborate the difference in the PCs’ output patterns, we analyzed the firing response in the function of spike adaptation, a phenomenon in which the inter-spike interval (ISI) gradually increases over time in response to a continuous current stimulus [24]. According to this analysis, the AF neurons exhibited higher slope values than IB and RS neurons, corroborating their higher adaptative nature (slope in AF = 0.12 ± 0.04, in IB = 0.03 ± 0.02, in RS = 0.03 ± 0.02; Fig 1E). Likewise, when plotting the instantaneous firing frequency (IFF) at different current intensities (120–360 pA) vs. the number of intervals (Fig 1F), RS neurons exhibited a near-to-steady firing frequency at distinct current intensities (Fig 1F1). On the other hand, the IB neurons initially exhibited a high firing frequency followed by a decay to a steady firing frequency (Fig 1F2), and the AF neurons showed a typical decreased firing frequency through stimulus, as a function of current magnitude (Fig 1F3).

### Layer V Pyramidal Cells in the Frontal Neocortex Exhibit Higher Firing Rates Relative to Temporal and Parietal Neurons of Patients with Drug-Resistant Epilepsy

Our previous results show that ≈66% of the recorded neurons belong to the RS phenotype. Therefore, we contrasted RS neurons’ passive and active electrophysiological properties by neocortex types, such as RMP, somatic input resistance (R_N_), and membrane time constant (τ). Fig 2A shows a representative neuron filled with biocytin, and Fig 2B1–B3 shows representative I–V curves from the different neocortices with the characteristic RS phenotype. Fig 2C summarizes the values of RMP between the three different neocortices (RMP in temporal neurons = -72.3 ± 1.35 mV, n = 6 cells/3 patients; in parietal neurons = - 73.4 ± 0.9 mV, n = 5 cells/3 patients; in frontal neurons = -73.13 ± 1.2 mV, n = 11 cells/4 patients). No statistical difference between groups was found in RMP. Next, we compared *R*_N_ and τ. However, neither *R*_N_ nor τ exhibited significant differences between neuronal groups (*R*_N_ in temporal neurons = 164.9 ± 17.8 MΩ; in parietal neurons = 163.9 ± 25.3 MΩ; in frontal neurons = 157.3 ± 8.2 MΩ; Fig 2D; τ in temporal neurons = 26.64 ± 3.35 ms; in parietal neurons = 32.7 ± 5.14 ms; in frontal neurons = 31.91 ± 3.14 ms; Fig 2E). We also determined the membrane capacitance of the cells belonging to the different cortices. However, we did not find differences between groups (membrane capacitance in temporal neurons = 166.4 ± 23.42 pF; in parietal neurons = 200.7 ± 11 pF; in frontal neurons = 214.5 ± 25.7 pF), suggesting that LV PCs across cortical regions in epileptic tissue have similar neuronal surfaces.

**Fig 2.**
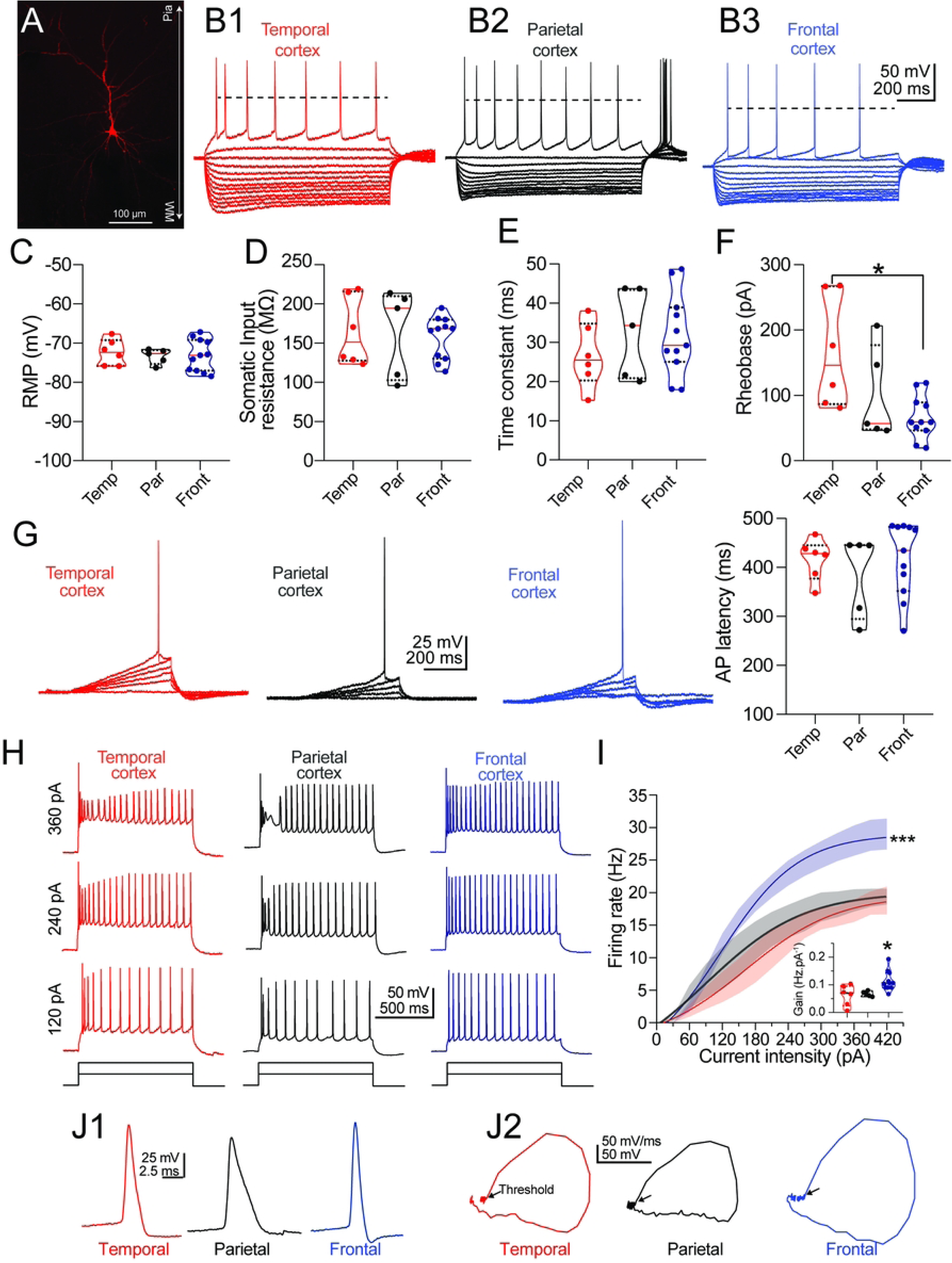
Passive membrane properties and neuronal firing in RS neurons from temporal, parietal, and frontal cortices of patients with drug-resistant epilepsy. **(A)** A typical PC filled with biocytin from the layer V temporal cortex. WM = white matter. (**B1– B3)** Representative I–V curves of RS PCs of the temporal cortex (B1, n = 6 cells/3 patients), parietal cortex (B2, n = 5 cells/3 patients), and frontal cortex (B3, n = 11 cells/4 patients). The I–V curves were elicited with current injections from -300 pA to 60 pA (30 pA steps). The horizontal dashed line represents the AP overshot. Passive and active parameters obtained from RS neurons: the violin plots contrast **(C)** the resting membrane potential (RMP), **(D)** the somatic input resistance (*R*_N_), and **(D)** the membrane time constant (τ) between PCs of the different cortices. **(F)** Violin plot summarizing the rheobase current required to elicit one AP by injecting a depolarizing current ramp, as illustrated in Fig 2G. Values for each neuronal group. * *p* < 0.05, one-way ANOVA followed Tukey’s test. **(G)** In the left panel, neuronal firing in PCs in response to the injection of a depolarizing ramp. The membrane potential was set at -70 mV. In the right panel, the violin plot shows the AP latency (in ms) elicited with rheobase current. **(H)** Representative firing traces from temporal (red), parietal (black), and frontal (blue) cortices to depolarizing current steps (1 s). **(I)** Line graph with frequency rate (Hz) – current of the different cortical neurons. Notice that PCs from the frontal cortex exhibited a higher frequency rate than temporal and parietal neurons. *** *p* < 0.001, mixed-effects ANOVA followed by Tukey’s test. Similarly, the comparison of neuronal gain (Hz pA^−1^), calculated as the slope of the linear portion (90–210 pA) of the frequency rate-current curve, corroborated higher excitability in frontal neurons (Fig 2I, violin plot inset). * *p* < 0.05, one-way ANOVA followed by Tukey’s test. **(J1)** Expanded AP traces elicited with rheobase current by cortex type and its representation as a phase plot **(J2)**. The phase plots were constructed by plotting the membrane voltage (mV) vs. its first derivative (mV/ms).

Next, we applied a depolarizing current ramp and determined the rheobase and the latency to elicit an AP in RS PCs from the distinct neocortex. Temporal and parietal RS neurons did not show differences (Fig 1F, left and middle panels); however, frontal neurons required lower current values to elicit an AP (rheobase in temporal neurons = 166 ± 35 pA; in parietal neurons = 101 ± 32.3 pA; in frontal neurons = 65.9 ± 10 pA; one-way ANOVA: F_(2, 19)_ = 5.405, Tukey’s test, *p* < 0.05; Fig 2F, right panel). On the other hand, we did not detect differences in latency to the AP at the rheobase current between groups (latency in temporal neurons = 416 ± 17.2 ms; in parietal neurons = 385 ± 37.6 ms; in frontal neurons = 416.6 ± 23 ms; Fig 2G).

Next, we compared the firing rate of RS PCs in response to increasing current injection (0– 390 pA). The analysis revealed that the maximal firing rates were similar for parietal and temporal neurons (18.2 ± 2.3 Hz vs. 18.4 ± 1.8 Hz, respectively; see black and blue traces, Fig 2H). In sharp contrast, RS PCs from the frontal neocortex exhibited a higher firing rate, reaching a maximal value of 28.82 ± 2.4 Hz (mixed-effects ANOVA, neocortex type effect: F_(2, 19)_ = 5.598, followed by Tukey’s test, *p* < 0.001 in frontal neurons vs. temporal and parietal neurons; Fig 2H–I). Consistent with these findings, when computing the neuronal gain, we corroborated the higher firing frequency of RS PCs from the frontal neocortex compared with the temporal and parietal neocortex (neuronal gain in temporal neurons = 0.062 ± 0.015 Hz.pA^−1^, in parietal neurons = 0.067 ± 0.004 Hz.pA^−1^, in frontal neurons = 0.114 ± 0.01 Hz.pA^−1^; one-way ANOVA, F_(2, 19)_ = 6.433, followed by Tukey’s test, *p* < 0.05; inset in Fig 2I).

Lastly, we examined the kinetic parameters of the AP waveform elicited at rheobase (Figs 2G and J1), including maximal depolarization slope (MDS), maximal repolarization slope (MRS), and half-width (H-W). Frontal PCs exhibited higher MDS values compared with temporal and parietal cells (MDS in temporal neurons = 179 ± 14.3 mV/ms; in parietal neurons = 184.3 ± 12.5 mV/ms; in frontal neurons = 247.2 ± 7.2 mV/ms; one-way ANOVA: F_(2, 19)_ = 15.07, followed by Tukey’s test, *p* < 0.05 in frontal vs. temporal and parietal PCs), suggesting higher global Na^+^ conductances in frontal PCs. By contrast, the comparison of the MRS, which reflects the activity of outward K^+^ conductances mediating the fast repolarization of the AP, showed no statistical differences between the neurons from the different neocortex (MRS in temporal neurons = -83.9 ± 15.1 mV/ms; in parietal neurons = -48.2 ± 0.9 mV/ms; in frontal neurons = -77.5 ± 8.3 mV/ms). Likewise, when comparing the H-W values between groups revealed that parietal neurons exhibit increased H-W values compared to temporal and frontal PCs (H-W in temporal neurons = 1.7 ± 0.3 ms; in parietal neurons = 2.7 ± 0.3 ms; in frontal neurons = 1.8 ± 0.1 ms; one-way ANOVA: F_(2, 19)_ = 4.796, followed by Tukey’s test, *p* < 0.05 in parietal vs. temporal and frontal PCs). The differences in kinetic parameters are depicted in the phase plots in Fig 2J2.

### Kinetic Properties of Neocortical fEPSPs and Synaptic Synchronization in Drug-Resistant Epilepsy

In parallel with the patch-clamp experiments, we also performed extracellular recordings in independent groups to explore the synaptic properties of the neocortical synapses. A stimulation electrode was placed in layer I/II, and the evoked response was recorded in the external region of layer V (Fig 1C). Parameters of the fEPSP, including amplitude, peak latency, and H-W, were analyzed (Fig 3A). The panels in Fig 3B1–B3 show representative fEPSPs of the temporal, parietal, and frontal neocortex. In the temporal neocortex (n = 8 slices/3 patients), the fEPSP was significantly larger compared with the parietal (n = 8 slices/3 patients) and frontal (n = 8 slices/4 patients) neocortical responses (fEPSP amplitude in temporal neocortex = 0.96 ± 0.04 mV; in parietal neocortex = 0.6 ± 0.07 mV; in frontal neocortex = 0.63 ± 0.07 mV; one-way ANOVA: F_(2, 21)_ = 9.767, Tukey’s test, *p* < 0.01 in temporal vs. parietal and frontal neocortex; Fig 3C). However, neither the peak latency nor the H-W of the fEPSPs was different between groups (peak latency in temporal neocortex = 1.67 ± 0.1 ms; in parietal neocortex = 1.75 ± 0.2 ms; in frontal neocortex = 2.06 ± 0.2 ms; Fig 3D; fEPSP H-W in temporal neocortex = 2.1 ± 0.2 ms; in parietal neocortex = 1.82 ± 0.2 ms; in frontal neocortex = 1.94 ± 0.23 ms; Fig 3E).

**Fig 3.**
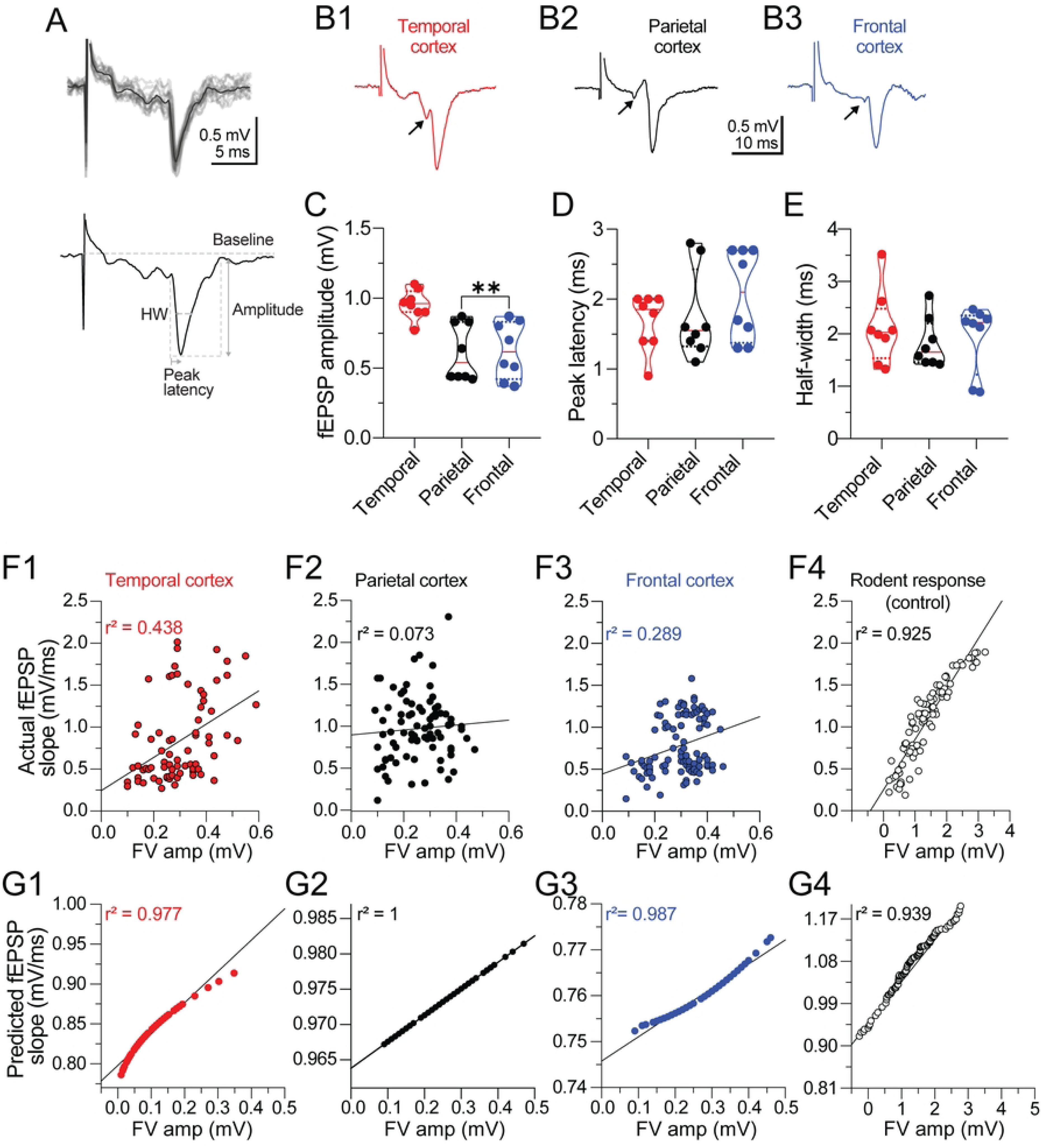
Kinetic profile of layer V cortical fEPSPs in patients with drug-resistant epilepsy. **(A)** The upper panel is a sequence of 10 consecutive fEPSPs recorded in layer V of the cortex in response to electrical stimulation in layer I/II (see schematic representation in Fig 1C). The individual traces are shown in gray, and the averaged response is in black. The lower panel shows the kinetic parameters analyzed from the fEPSP waveforms, including fEPSP amplitude (mV), latency to the peak of the fEPSP (peak latency, ms), and fEPSP half-width (HW, ms). **(B)** Representative fEPSP traces from **(B1)** temporal, **(B2)** parietal, and **(B3)** frontal neocortex. Each trace was averaged from five continuous sweeps acquired at 0.067 Hz. The arrowhead preceding the fEPSP is the presynaptic fiber volley. In the lower panel, violin plots summarize the kinetic parameters of **(C)** the fEPSP amplitude, **(D)** peak latency, and **(E)** fEPSP H-W, obtained from the three cortices. Notice that fEPSP amplitudes from parietal and frontal slices are smaller compared to temporal slices. n = 8 slices/3 patients for temporal and parietal slices; n = 8 slices/4 patients for frontal slices. \**p* < 0.05, one-way ANOVA followed by Tukey’s test. **(F1–F3)** Scatter plot showing the individual relationship between presynaptic fiber volley vs. actual fEPSP amplitude from the three cortical areas and the resulting Pearson’s correlation coefficient. Each dot within the plot represents an actual sweep that included FV and fEPSP. **(F4)** Scatter plot showing the same relationship in rodents. Notice the differences in Pearson’s *r*^2^ coefficient between DRE and control responses. **(G1–G3)** Scatter plots showing the theoretical consistency and proportionality of the predicted FV vs. fEPSP response for each cortical area according to the least square methods. Under theoretical conditions, cortical responses should be linearly correlated, as occurs in the **(G4)** scatter plot of control responses acquired from rodent brain.

During the acquisition of the fEPSPs, the amplitude of the fiber volley (FV or presynaptic action potentials) and the fEPSP exhibited amplitude variability (i.e., small FVs and larger fEPSPs or vice versa), despite maintaining constant current and stimulation frequency (0.067 Hz). Previous studies in the rodent’s central synapses indicate that the FV–fEPSP relationship is linear, and pathological conditions alter this relationship [30,31]. Therefore, to explore a possible desynchronization between presynaptic excitability and postsynaptic response, the individual trial-to-trial values of FV vs. its respective fEPSP slopes were correlated for each human neocortex (Fig 3F1–F3). We also plotted the relationship of FV vs. fEPSPs from a control rodent’s acute slice for comparative purposes. In the DRE neocortical tissue, Pearson’s *r*^2^ coefficients suggested a lack of high correlation between FV vs. fEPSP (temporal cortex *r*^2^ = 0.438; parietal *r*^2^ = 0.073; frontal *r*^2^ = 0.289; Fig 3F1–F3). In sharp contrast, *r*^2^ in the rodent response was a high linear correlation (*r*^2^ = 925; Fig 3F4). We applied the least squares method to the FV-fEPSP data from each cortical area to further substantiate this possible synaptic imbalance. In response to the adjustments obtained from the least-square methods, Fig 3G1–G3 shows the theoretical consistency and proportionality of the predicted FV vs. fEPSP response for each cortical area (temporal cortex *r*^2^ = 0.9541, parietal *r*^2^ = 0.9745, and frontal *r*^2^ = 1). These data indicate a foreseeable linear relationship between FV amplitude and fEPSP, where increases in FV amplitude correspond to linear increases in fEPSP amplitude. However, in pathological conditions such as DRE, this proportionality is disrupted.

### Neocortical Synapses From Patients With Drug-Resistant Epilepsy Exhibit Loss of Short-Term Depression

A distinctive property of the neocortical excitatory synapses of rodents is its low probability of neurotransmitter release [32], a phenomenon that governs glutamatergic transmission’s paired-pulse ratio (PPR). However, PPR has been barely explored in the human neocortical synapses. Therefore, the next series of experiments aimed to determine the PPR of the glutamatergic transmission at the layer I/II–V PCs synapses using an inter-stimulus interval of 60 ms. Figs 4A and 4B show the mild or spare PPR facilitation (PPF) found in the three neocortical areas. In the temporal neocortex, the PPR was 1.03 ± 0.06 (n = 6 slices/3 patients), while in the parietal and frontal neocortex, PPR was 1.07 ± 0.05 (n = 7 slices/3 patients) and 1.15 ± 0.11 (n = 7 slices/4 patients), respectively (Fig 4B).

**Fig 4.**
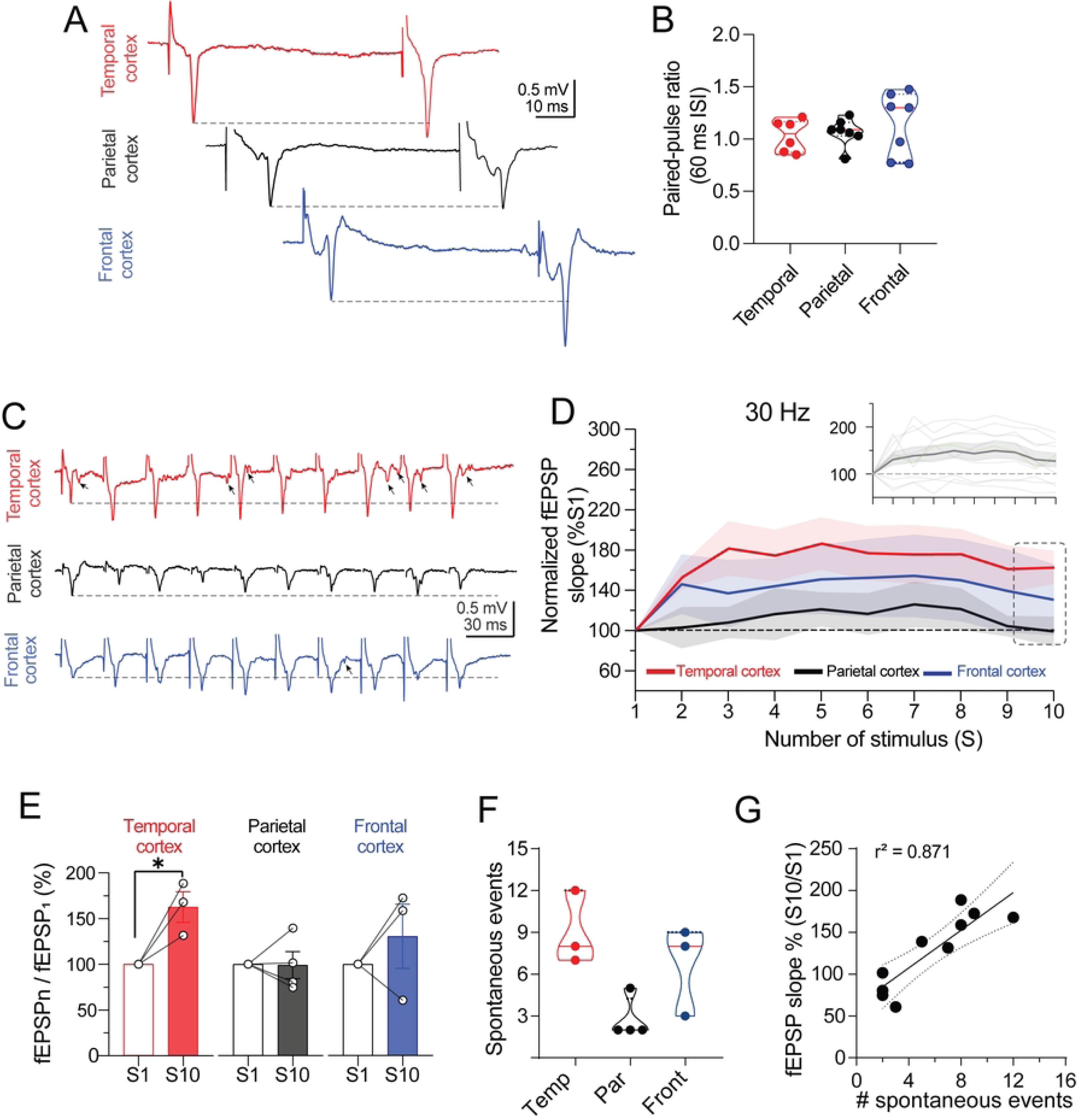
LI/II–V cortical synapses from patients with drug-resistant epilepsy lack frequency-dependent short-term depression. **(A)** Representative traces of paired stimulation (60 ms) from the temporal, parietal, and frontal neocortex. **(B)** Violin plot contrasting the paired-pulse ratio (PPR) between cortical groups. On average, the PPR showed spare facilitation in each cortical synapse. n = 6 slices/3 patients for the temporal slices; n = 7 slices/3 patients for the parietal slices; n = 7 slices/4 patients for frontal slices. **(C)** Representative traces of evoked fEPSPs in response to a train of 10 pulses elicited at 30 Hz in the temporal, parietal, and frontal cortices. The horizontal dashed line represents the facilitation level. **(D)** Line plot of each evoked synaptic response recorded in layer V during the stimulation trains delivered in LI/II at 30 Hz. At the individual level, the recorded trains exhibited both synaptic depression and facilitation, and the average response was biased toward synaptic facilitation (green traces, inset panel D). When sorting the slices by cortex type, the temporal and frontal synapses exhibited synaptic facilitation, whereas parietal responses did not show any facilitation. **(E)** The bar and line graphs show that only temporal synapses exhibited a significant facilitation level (fEPSP slope % of S10 vs. S1). \**p* < 0.05, ratio-paired t-test. n = 3 slices/3 patients for temporal and parietal slices; n = 4 slices/3 patients for parietal slices. **(F)** Violin plot summarizing the number of spontaneous synaptic events recorded during 30 Hz train stimulation across different neocortical regions. **(G)** Pearsońs correlation analysis between the number of spontaneous synaptic events and the magnitude of the tenth synaptic response during 30 Hz train stimulation.

According to previous studies, repetitive synaptic stimulation within the gamma range triggers short-term depression (STD) in rodent and human neocortices [15,32,33]. Therefore, we examined the epileptic neocortex’s response to trains of ten stimuli (S) at 30 Hz. To avoid synaptic saturation during the experimental trials, we set up the magnitude of the applied current to evoke ≈50% of the maximal amplitude of the fEPSP in each slice [25]. These responses exhibited a dominant synaptic facilitation independent of the neocortex type (inset Fig 4D). When distinguished by neocortical area (Fig 4D), temporal and frontal neocortex exhibited sustained facilitation in response to a 30 Hz train (red and blue lines, respectively), whereas the parietal neocortex showed unfacilitated responses (black line). The temporal neocortex was the only region with a significant synaptic facilitation in response to 10 Hz train (fEPSP_s10_/fEPSP_s1_ = 162.7 ± 16.7% of S1; ratio-paired t-test, t_(2)_ = 4.48, *p* < 0.05; Fig 4E).

Additionally, we observed that during train stimulation, spontaneous synaptic events emerged following evoked synaptic responses, with latencies exceeding 10 ms after electrical stimuli (see arrows in Fig 4C). The number of these synaptic events tended to be higher in slices from the temporal neocortex than in those from the parietal or frontal neocortex (Fig 4F). Interestingly, we found a strong positive correlation between the number of spontaneous synaptic events and the magnitude of the last synaptic response during the 30 Hz train (Pearson correlation, r² = 0.871; Fig 4G). These data suggest that frequency-dependent synaptic facilitation or the loss of STD in epileptic neocortex favors excitatory synaptic activity.

### Morphometric Analysis of Layer V Pyramidal Cells

Because the patch pipettes included biocytin (see methods), post hoc analyses were used to reveal details of PCs’ dendritic organization. The analysis was performed in four RS PCs obtained from the parietal and frontal neocortex. The cells exhibited similar morphometric and electrophysiological properties (Fig 5A–B). Seven additional PCs exhibited sparse biocytin labeling, restricted to the somata or random sections of the dendritic ramifications. These cells were discarded.

**Fig 5.**
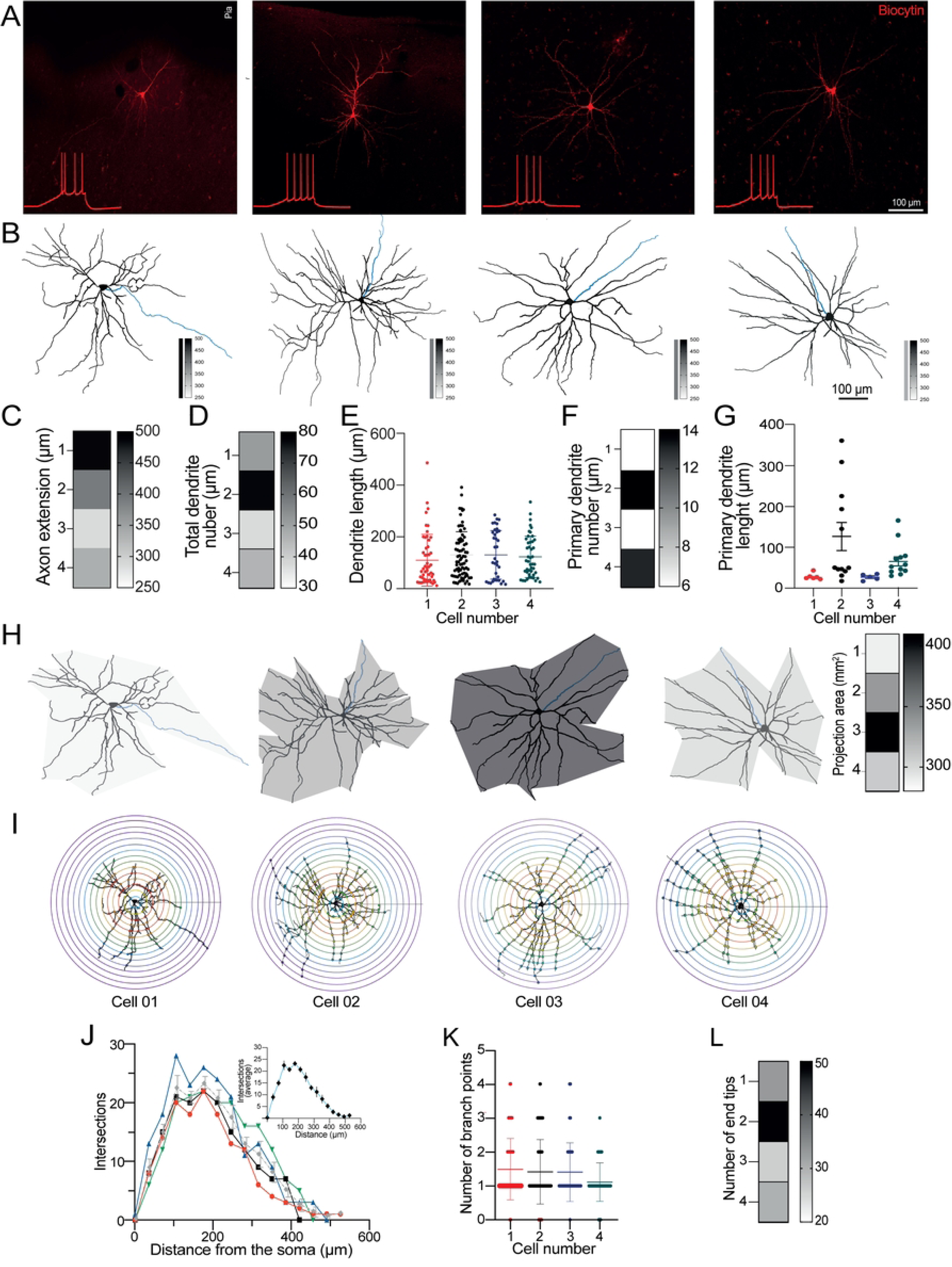
Morphological analysis and dendritic complexity of layer V pyramidal neurons with regular spiking pattern. **(A)** Immunofluorescence photomicrographs of PCs filled with biocytin via the patch pipette. The composite images result from an average of at least 40 Z-stacks. Inset, firing response of the labeled neurons. **(B)** Post hoc digital reconstructions were performed with the Simple NeuroTracing toolbox of ImageJ. Dendritic arborizations are shown in black, and the axon in blue. The inset next to each reconstruction shows each neuron’s axon extension heatmap. **(C)** Heatmap of the axon extension, ranging from white to black, with black indicating the most extensive axon length. **(D)** The heat map of the total dendrites shows that black indicates the highest number. **(E)** Scatter plot with the total dendrite length for each neuron. **(F)** Heatmap of the number of primary dendrites for each neuron; black indicates the highest number of primary dendrites. **(G)** Scatter plot contrasting the length of primary dendrites for each neuron. **(H)** The total area extension for each neuron and heatmap is black, which indicates the largest extension area. Each color in the area corresponds to the heatmap. **(I)** Sholl analysis was performed with the concentric spheres with a 35 µM diameter. **(J)** Individual analysis showing the dendritic complexity of each PC. The inset graph represents the average response ± SEM. **(K)** Scatter plot summarizing the number of branch points of each neuron. **(L)** The heat map shows dendritic branching end tips, with black indicating the highest number.

The axon’s initial segment was visually identified for all the reconstructed cells. The axon stemmed from the soma base and had a thin appearance with a mean extension of 54.05 ± 15.4 μm (blue traces in Fig 5B; total quantification in Fig 5C). Next, the total number of dendrites and the measurement of their total length had comparable values between cells (Figs 5D and 5E; see also Table 2). Likewise, the number and the primary dendritic length exhibited similar values (Figs 5F and 5G; Table 2). Furthermore, despite variations in dendritic distribution among cells, their surface areas were similar (projection area for cell 1 = 287.26 μm²; for cell 2 = 336.43 μm²; for cell 3 = 408.71 μm²; for cell 4 = 307.59 μm²; Fig 5H).

**Table 2.** Morphological measurements and Sholl Analysis of Layer V pyramidal cells from patients with drug-resistant epilepsy. The dendritic measurements were performed in four RS PCs obtained from the parietal and frontal neocortices.

| **Morphological measurements** |  |  |  |  |
| --- | --- | --- | --- | --- |
| # Cell | number of total dendrites | total length of the dendrites | number of total primary dendrites | primary dendritic length |
| Cell 01 Parietal | 51 | 107.9 ± 14.1 µm | 6 | 24.44 ± 4.69 µm |
| Cell 02 Parietal | 77 | 129.7 ± 10.3 µm | 14 | 117 ± 33.05 µm |
| Cell 03 Frontal | 37 | 130.4 ± 13.4 µm | 6 | 23 ± 4.80 µm |
| Cell 04 Frontal | 46 | 122.7 ± 11.8 µm | 13 | 60.64 ± 11.26 µm |
| **Sholl Analysis** |  |  |  |  |
| # Cell | Number of branching points |  | Branching end tips |  |
| Cell 01 Parietal | 1.52 ± 0.12 |  | 33 |  |
| Cell 02 Parietal | 1.42 ± 0.14 |  | 49 |  |
| Cell 03 Frontal | 1.46 ± 0.18 |  | 26 |  |
| Cell 04 Frontal | 1.19 ± 0.12 |  | 30 |  |

Next, the dendritic tree branching patterns and complexity were evaluated with a Sholl analysis (Figs 5I and 5J). The quantification of the branching points showed no statistical differences between the cells (Fig 5K; Table 2), and similar results were found for branching end tips (Fig 5L; Table 2). Surprisingly, the relation between the number of branch points and end tips was appropriate (r = 0.99). According to these morphometric analyses, our results indicate that neocortical tissue from patients with DRE has dysmorphic neurons exhibiting dendritic alterations, such as simplified branching, reduced arborization, and decreased complexity.

## Discussion

This study summarizes a series of electrophysiological and morphological properties of layer V PCs from three neocortical regions surgically resected from patients with DRE. At the cellular level, we used patch-clamp recordings to classify PCs according to their firing pattern and to determine membrane subthreshold and suprathreshold properties. At the circuit level, extracellular recordings were used to evaluate synaptic strength, synchronization between presynaptic volleys and excitatory postsynaptic potentials, and frequency-dependent short-term plasticity within the gamma range. Lastly, we also performed post hoc neuronal reconstructions and analyses that revealed morphometric features of layer V PCs. Significantly, PCs from deep cortical layers such as layer V have been identified as a major source of epileptiform discharges in both animal models and human patients [19,21].

### Membrane Properties of PCs

Despite extensive knowledge of the neuronal physiology of rodent neurons, our understanding of the human neocortex’s intrinsic electrophysiological properties is still limited. Moreover, a recent study showed that some intrinsic properties, such as somatic R*_N_*, rheobase current to evoke an AP, currents mediated by HCN channels, and the firing of PCs differ between rodent and human neocortices [16].

Given the inherent challenge of obtaining healthy human brain tissue for comparative analysis [11], in most studies, tissue resected from areas away from epileptic zones or deep brain tumors is regarded ‘normal’ or ‘moderately less affected’ by neuropathologists [16,34]. The neurons examined in our study exhibited hyperpolarized RMP, minimal SSA, and firing outputs resembling those observed in rodent and human neocortices under normal or near-physiological conditions [16,35,36]. However, the homogeneity of the passive properties measurements with the patch-clamp recordings suggest that a subset of ion channels active near the RMP of these cells did not experience significant changes, thereby maintaining the ionic distribution across the cellular membrane. According to this idea, the higher excitability observed in frontal PCs, compared to parietal and temporal PCs, is more likely due to a higher density of Na^+^ channels (i.e., higher MDS values) rather than to changes in passive properties.

Another important consideration is that the electrophysiological measurements may vary depending on the age range of the individuals from whom the recordings were obtained. In the temporal neocortex, the neurons from patients aged 21 to 27 years exhibited a more hyperpolarized RMP (mean RMP ≈ -72 mV) compared to those neurons from normal neocortical tissue of patients aged 19 to 63 years (mean RMP ≈ -65 mV) [22] and patients aged 23 to 30 years (mean RMP ≈ -68 mV) (Allen Brain Atlas database). It is necessary to analyze more neurons to determine if the RMP is stable throughout the lifespan or if its value fluctuates with neurological diseases. Despite the noticeable age fluctuation among the examined neocortical areas (2–30 years) as a limitation of our study, the homogeneity of age within the parietal neocortex group (2–7 years, childhood) did not suggest possible differences in the intrinsic cellular properties of PCs compared to the temporal neocortex group, which exhibited a similar age homogeneity (21–27 years, adulthood). However, analyzing a larger number of cells in future studies should help clarify this observation.

### Changes in the Synaptic Transmission and Short-Term Plasticity of the Neocortical Synapses

Within neocortical synaptic networks, the postsynaptic firing is influenced, among other factors, by the dynamics of presynaptic release and the short-term plasticity of synaptic transmission [33,37]. Consistent with findings in the rodent neocortex, in near-physiological conditions, frequency-dependent STD modulates input-efficient information transfer among distinct levels of the human PCs of different neocortical layers [15].

Likewise, STD balances neocortical activity during prolonged synaptic stimulation [33], positing frequency-dependent STD as a key cellular mechanism for regulating the propagation of electrical activity across the neocortical layers. With that in mind, our extracellular recordings uncovered a loss of synaptic depression in response to gamma range stimulation of the superficial neocortical layers I/II; even moderate synaptic facilitation was observed in the temporal neocortex (Fig 4C–D). Consequently, the loss of synaptic depression or gain of synaptic facilitation, both frequency-dependent events, may be a potential mechanism that favors network hyperexcitability during seizure activity by amplification of neuronal activity (e.g., summation of fEPSPs during high-frequency activity). This possibility arises from the observation that synaptic facilitation during spike activity transiently increases neurotransmitter release probability, thereby enhancing the postsynaptic excitatory conductances [38]. Moreover, we observed that the loss of frequency-dependent STD was highly correlated with a higher excitatory synaptic activity. Finally, a potential explanation for the loss of frequency-dependent STD is the decreased expression of HCN1 channels, given that the inactivation or blockade of these channels not only increases the temporal summation of synaptic events but also promotes epileptiform discharges [39,40]. If this is true, promoting the activity of HCN1 channels in certain types of epilepsy, but see [41], may restore the frequency-dependent STD and consequently prevent or reduce the cortical spread of pathological electrical activity during seizures, as these channels preferentially regulate dendritic excitability and epileptogenesis [42,43].

In the human cortex, presynaptic release probability has been barely explored, yet, it is documented that layer II/III synapses of the temporal neocortex from non-epileptic tissue exhibit paired-pulse depression [15]. This is consistent with previous studies using dual patch-clamp recordings from PCs of the rat neocortex [33]. In this sense, we found that paired stimulation (60 ms) in all the neocortical synapses did not trigger depression; contrary to the expected, a mild paired-pulse facilitation was observed. This apparent polarity shift suggests a change in the neurotransmitter release process from high to low release probability [44]. We hypothesize that the reduced synaptic depression (i.e., PPF) in the temporal neocortex increases the excitability during high-frequency activity, such as is observed during seizures. A similar phenomenon has been previously documented in murine neurons [45].

### Morphometric Alterations of Layer V Neocortical Neurons

Because biocytin was included in the patch-clamp pipettes, the tracer diffused throughout the intracellular compartments while the neurons were recorded. Post hoc treatments and immunofluorescence imaging allowed us to trace neuronal morphology and, in some cases, detect neuronal coupling (not shown). The morphometric and Sholl analyses showed reduced dendritic branching and complexity. This reduced dendritic complexity is expected to impair synaptic transmission in individuals with epilepsy [46]. Likewise, the apical and basal dendrites exhibited irregular dendritic arborizations despite following a linear trajectory. These alterations may contribute to the dysregulation of transmission and synaptic plasticity in the epileptic brain [5,6,47], as observed in the synaptic transmission experiments (Figs 3 and 4). Finally, the Sholl analyses showed decreased bifurcations and complexity and simplification of dendritic branching patterns. Given that chronic seizures promote retraction of dendrites and loss of dendritic spines [48], the decreased dendritic branching observed in this study can be another consequence of DRE.

It is important to mention that the major limitation of our morphometric analyses is the reduced number of analyzed cells. Despite this limitation, each cell was electrophysiologically characterized and shared similar electrophysiological and morphological characteristics. In the near future, experiments with a larger population of human neurons from DRE will help to substantiate or rule out the present observations.

## Conclusion

Our findings suggest that layer V neocortical PCs from patients with drug-resistant epilepsy are not inherently hyperexcitable at the somatic level. Instead, their firing frequency is comparable to unaffected human tissue [16]. At the synaptic level, however, alterations in the PPR, desynchronization between presynaptic excitability and postsynaptic response, and a loss of frequency-dependent STD may significantly contribute to the hyperexcitable state observed during seizure activity. Future studies should aim to analyze the individual contributions of cellular elements within neuronal compartments to clarify their specific roles in the hyperexcitability underlying seizure activity.

## Acknowledgments

This work was supported by Conahcyt to LR A3-S-26782; Cinvestav-IPN to EJG. Additional funding was provided by doctoral fellowships 725800 (LAM) by Conahcyt. We would like to acknowledge R. Olvera for early preliminary experiments.

## Author Contributions

**Conceptualization:** Luis Alfredo Márquez, Emilio J. Galván

**Data curation:** Luis A. Márquez, Emilio J. Galván

**Formal analysis:** Luis A. Márquez, Christopher Martínez-Aguirre, Estefanía Gutierrez-Castañeda, Ernesto Griego, Isabel Sollozo-Dupont

**Funding acquisition:** Mario Alonso-Vanegas, Luisa Rocha,

**Investigation:** Luis A. Márquez, Christopher Martínez-Aguirre; Emilio J. Galván

**Methodology:** Luis A. Márquez, Christopher Martínez-Aguirre; Isabel Sollozo-Dupont, Emilio J. Galván

**Project administration:** Luisa Rocha, Emilio J. Galván

**Resources:** Mario Alonso-Vanegas, Luisa Rocha, Emilio J. Galván

**Validation:** Emilio J. Galván

**Writing - original draft:** Luis A. Márquez, Estefanía Gutierrez-Castañeda, Isabel Sollozo-Dupont, Emilio J. Galván

**Writing – review & editing:** Emilio J. Galván.

## References

1. McNamara JO. Emerging insights into the genesis of epilepsy. Nature. 1999;399: A15–A22. doi:10.1038/399a015.

2. Scharfman HE. The neurobiology of epilepsy. Current Neurology and Neuroscience Reports. Springer; 2007. pp. 348–354. doi:10.1007/s11910-007-0053-z.

3. Stöber TM, Batulin D, Triesch J, Narayanan R, Jedlicka P. Degeneracy in epilepsy: multiple routes to hyperexcitable brain circuits and their repair. Commun Biol. 2023;6: 1–16. doi:10.1038/s42003-023-04823-0.

4. Bernard C, Anderson A, Becker A, Poolos NP, Deck H, Johnston D. Acquired dendritic channelopathy in temporal lobe epilepsy. Science (80-). 2004;305: 532–535. doi:10.1126/science.1097065.

5. Freiman TM, Eismann-Schweimler J, Frotscher M. Granule cell dispersion in temporal lobe epilepsy is associated with changes in dendritic orientation and spine distribution. Exp Neurol. 2011;229. doi:10.1016/j.expneurol.2011.02.017.

6. Masala N, Pofahl M, Haubrich AN, Sameen Islam KU, Nikbakht N, Pasdarnavab M, et al. Targeting aberrant dendritic integration to treat cognitive comorbidities of epilepsy. Brain. 2023;146. doi:10.1093/brain/awac455.

7. Tóth K, Hofer KT, Kandrács Á, Entz L, Bagó A, Erőss L, et al. Hyperexcitability of the network contributes to synchronization processes in the human epileptic neocortex. J Physiol. 2018;596: 317–342. doi:10.1113/JP275413-

8. Rich S, Moradi Chameh H, Lefebvre J, Valiante TA. Loss of neuronal heterogeneity in epileptogenic human tissue impairs network resilience to sudden changes in synchrony. Cell Rep. 2022;39: 110863. doi:10.1016/j.celrep.2022.110863.

9. Dominguez LG, Wennberg RA, Gaetz W, Cheyne D, Snead OC, Perez Velazquez JL. Enhanced synchrony in epileptiform activity? Local versus distant phase synchronization in generalized seizures. J Neurosci. 2005;25: 8077–8084. doi:10.1523/JNEUROSCI.1046-05.2005.

10. Farrell JS, Nguyen QA, Soltesz I. Resolving the Micro-Macro Disconnect to Address Core Features of Seizure Networks. Neuron. 2019;101: 1016–1028. doi:10.1016/J.NEURON.2019.01.043.

11. Moradi Chameh H, Falby M, Movahed M, Arbabi K, Rich S, Zhang L, et al. Distinctive biophysical features of human cell-types: insights from studies of neurosurgically resected brain tissue. Front Synaptic Neurosci. 2023;15: 1250834. doi:10.3389/FNSYN.2023.1250834.

12. Larkum ME, Nevian T, Sandier M, Polsky A, Schiller J. Synaptic integration in tuft dendrites of layer 5 pyramidal neurons: A new unifying principle. Science (80-). 2009;325: 756–760. doi:10.1126/SCIENCE.1171958.

13. Shai AS, Anastassiou CA, Larkum ME, Koch C. Physiology of Layer 5 Pyramidal Neurons in Mouse Primary Visual Cortex: Coincidence Detection through Bursting. PLOS Comput Biol. 2015;11: e1004090. doi:10.1371/JOURNAL.PCBI.1004090.

14. Wilbers R, Metodieva VD, Duverdin S, Heyer DB, Galakhova AA, Mertens EJ, et al. Human voltage-gated Na+ and K+ channel properties underlie sustained fast AP signaling. Sci Adv. 2023;9: eade3300. doi:10.1126/SCIADV.ADE3300.

15. Testa-Silva G, Verhoog MB, Linaro D, de Kock CPJ, Baayen JC, Meredith RM, et al. High Bandwidth Synaptic Communication and Frequency Tracking in Human Neocortex. PLOS Biol. 2014;12: e1002007. doi:10.1371/JOURNAL.PBIO.1002007.

16. Beaulieu-Laroche L, Toloza EHS, van der Goes MS, Lafourcade M, Barnagian D, Williams ZM, et al. Enhanced Dendritic Compartmentalization in Human Cortical Neurons. Cell. 2018;175: 643–651.e14. doi:10.1016/j.cell.2018.08.045-

17. Mohan H, Verhoog MB, Doreswamy KK, Eyal G, Aardse R, Lodder BN, et al. Dendritic and Axonal Architecture of Individual Pyramidal Neurons across Layers of Adult Human Neocortex. Cereb Cortex. 2015;25: 4839–4853. doi:10.1093/CERCOR/BHV188.

18. Ofer N, Benavides-Piccione R, Defelipe J, Yuste R. Structural Analysis of Human and Mouse Dendritic Spines Reveals a Morphological Continuum and Differences across Ages and Species. eNeuro. 2022;9: 1–12. doi:10.1523/ENEURO.0039-22.2022.

19. Hoffman SN, Salin PA, Prince DA. Chronic neocortical epileptogenesis in vitro. J Neurophysiol. 1994;71: 1762–1773. doi:10.1152/JN.1994.71.5.1762.

20. Telfeian AE, Connors BW. Layer-Specific Pathways for the Horizontal Propagation of Epileptiform Discharges in Neocortex. Epilepsia. 1998;39: 700–708. doi:10.1111/J.1528-1157.1998.TB01154.X.

21. Bourdillon P, Ren L, Halgren M, Paulk AC, Salami P, Ulbert I, et al. Differential cortical layer engagement during seizure initiation and spread in humans. Nat Commun. 2024;15: 1–13. doi:10.1038/s41467-024-48746-8.

22. Moradi Chameh H, Rich S, Wang L, Chen F-D, Zhang L, Carlen PL, et al. Diversity amongst human cortical pyramidal neurons revealed via their sag currents and frequency preferences. Nat Commun. 2021;12: 2497. doi:10.1038/s41467-021-22741-9.

23. Planert H, Xaver Mittermaier F, Grosser S, Fidzinski P, Schneider UC, Radbruch H, et al. Cellular and Synaptic Diversity of Layer 2-3 Pyramidal Neurons in Human Individuals. bioRxiv. 2023; 2021.11.08.467668. doi:10.1101/2021.11.08.467668.

24. Hemond P, Epstein D, Boley A, Migliore M, Ascoli GA, Jaffe DB. Distinct classes of pyramidal cells exhibit mutually exclusive firing patterns in hippocampal area CA3b. Hippocampus. 2008;18: 411–424. doi:10.1002/HIPO.20404.

25. Márquez LA, López Rubalcava C, Galván EJ. Postnatal hypofunction of N-methyl-D-aspartate receptors alters perforant path synaptic plasticity and filtering and impairs dentate gyrus-mediated spatial discrimination. Br J Pharmacol. 2024;181: 2701–2724. doi:10.1111/BPH.16375.

26. Cruz SL, Torres-Flores M, Galván EJ. Repeated toluene exposure alters the synaptic transmission of layer 5 medial prefrontal cortex. Neurotoxicol Teratol. 2019;73: 9–14. doi:10.1016/J.NTT.2019.02.002.

27. Griego E, Segura-Villalobos D, Lamas M, Galván EJ. Maternal immune activation increases excitability via downregulation of A-type potassium channels and reduces dendritic complexity of hippocampal neurons of the offspring. Brain Behav Immun. 2022;105: 67–81. doi:10.1016/J.BBI.2022.07.005.

28. Ferreira TA, Blackman A V., Oyrer J, Jayabal S, Chung AJ, Watt AJ, et al. Neuronal morphometry directly from bitmap images. Nat Methods 2014 1110. 2014;11: 982–984. doi:10.1038/nmeth.3125.

29. Gutierrez-Castañeda NE, Martínez-Rojas VA, Ochoa-de la Paz LD, Galván EJ. The bidirectional role of GABAA and GABAB receptors during the differentiation process of neural precursor cells of the subventricular zone. PLoS One. 2024;19: e0305853. doi:10.1371/JOURNAL.PONE.0305853.

30. Barnes CA, McNaughton BL. Physiological compensation for loss of afferent synapses in rat hippocampal granule cells during senescence. J Physiol. 1980;309: 473–485. doi:10.1113/JPHYSIOL.1980.SP01352.

31. Villanueva-Castillo C, Tecuatl C, Herrera-López G, Galván EJ. Aging-related impairments of hippocampal mossy fibers synapses on CA3 pyramidal cells. Neurobiol Aging. 2017;49: 119–137. doi:10.1016/J.NEUROBIOLAGING.2016.09.010.

32. Crabtree GW, Sun Z, Kvajo M, Broek JAC, Fénelon K, McKellar H, et al. Alteration of Neuronal Excitability and Short-Term Synaptic Plasticity in the Prefrontal Cortex of a Mouse Model of Mental Illness. J Neurosci. 2017;37: 4158–4180. doi:10.1523/JNEUROSCI.4345-15.2017.

33. Galarreta M, Hestrin S. Frequency-dependent synaptic depression and the balance of excitation and inhibition in the neocortex. Nat Neurosci. 1998;1: 587–594. doi:10.1038/2822.

34. Testa-Silva G, Verhoog MB, Goriounova NA, Loebel A, Hjorth JJJ, Baayen JC, et al. Human synapses show a wide temporal window for spike-timing-dependent plasticity. Front Synaptic Neurosci. 2010;2: 1388. doi:10.3389/FNSYN.2010.00012/BIBTEX.

35. Armenta-Resendiz M, Cruz SL, Galván EJ. Repeated toluene exposure increases the excitability of layer 5 pyramidal neurons in the prefrontal cortex of adolescent rats. Neurotoxicol Teratol. 2018;68: 27–35. doi:10.1016/J.NTT.2018.04.006.

36. Griego E, Santiago-Jiménez G, Galván EJ. Systemic administration of lipopolysaccharide induces hyperexcitability of prelimbic neurons via modulation of sodium and potassium currents. Neurotoxicology. 2022;91: 128–139. doi:10.1016/J.NEURO.2022.05.010.

37. Brenowitz S, Trussell LO. Minimizing Synaptic Depression by Control of Release Probability. J Neurosci. 2001;21: 1857–1867. doi:10.1523/JNEUROSCI.21-06-01857.2001.

38. Tauffer L, Kumar A. Short-Term Synaptic Plasticity Makes Neurons Sensitive to the Distribution of Presynaptic Population Firing Rates. eNeuro. 2021;8: 1–23. doi:10.1523/ENEURO.0297-20.2021.

39. Thuault SJ, Malleret G, Constantinople CM, Nicholls R, Chen I, Zhu J, et al. Prefrontal Cortex HCN1 Channels Enable Intrinsic Persistent Neural Firing and Executive Memory Function. J Neurosci. 2013;33: 13583–13599. doi:10.1523/JNEUROSCI.2427-12.2013.

40. Albertson AJ, Williams SB, Hablitz JJ. Regulation of epileptiform discharges in rat neocortex by HCN channels. J Neurophysiol. 2013;110: 1733–1743. doi:10.1152/JN.00955.2012.

41. Bleakley LE, Reid CA. HCN1 epilepsy: From genetics and mechanisms to precision therapies. J Neurochem. 2024;168: 3891–3910. doi:10.1111/JNC.15928.

42. Poolos NP, Migliore M, Johnston D. Pharmacological upregulation of h-channels reduces the excitability of pyramidal neuron dendrites. Nat Neurosci . 2002;5: 767–774. doi:10.1038/nn891.

43. Shah MM, Anderson AE, Leung V, Lin X, Johnston D. Seizure-induced plasticity of h channels in entorhinal cortical layer III pyramidal neurons. Neuron. 2004;44: 495–508. doi:10.1016/J.NEURON.2004.10.011.

44. Regehr WG. Short-Term Presynaptic Plasticity. Cold Spring Harb Perspect Biol. 2012;4: a005702. doi:10.1101/CSHPERSPECT.A005702.

45. Lou X, Fan F, Messa M, Raimondi A, Wu Y, Looger LL, et al. Reduced release probability prevents vesicle depletion and transmission failure at dynamin mutant synapses. Proc Natl Acad Sci U S A. 2012;109: E515–E523. doi:10.1073/PNAS.1121626109.

46. Rossini L, de Santis D, Mauceri RR, Tesoriero C, Bentivoglio M, Maderna E, et al. Dendritic pathology, spine loss and synaptic reorganization in human cortex from epilepsy patients. Brain. 2021;144. doi:10.1093/brain/awaa387.

47. Beck H, Goussakov I V., Lie A, Helmstaedter C, Elger CE. Synaptic Plasticity in the Human Dentate Gyrus. J Neurosci. 2000;20: 7080–7086. doi:10.1523/JNEUROSCI.20-18-07080.2000.

48. Runge K, Cardoso C, de Chevigny A. Dendritic Spine Plasticity: Function and Mechanisms. Frontiers in Synaptic Neuroscience. 2020. doi:10.3389/fnsyn.2020.00036.

